# A machine learning algorithm predicts molecular subtypes in pancreatic ductal adenocarcinoma with differential response to gemcitabine-based versus FOLFIRINOX chemotherapy

**DOI:** 10.1101/664540

**Authors:** Georgios Kaissis, Sebastian Ziegelmayer, Fabian Lohöfer, Katja Steiger, Hana Algül, Alexander Muckenhuber, Hsi-Yu Yen, Ernst Rummeny, Helmut Friess, Roland Schmid, Wilko Weichert, Jens T. Siveke, Rickmer Braren

## Abstract

**Purpose:** Development of a supervised machine-learning model capable of predicting clinically relevant molecular subtypes of pancreatic ductal adenocarcinoma (PDAC) from diffusion-weighted-imaging-derived radiomic features.

**Methods:** The retrospective observational study assessed 55 surgical PDAC patients. Molecular subtypes were defined by immunohistochemical staining of KRT81. Tumors were manually segmented and 1606 radiomic features were extracted with *PyRadiomics*. A gradient-boosted-tree algorithm (XGBoost) was trained on 70% of the patients (N=28) and tested on 30% (N=17) to predict KRT81+ vs. KRT81-tumor subtypes. The average sensitivity, specificity and ROC-AUC value were calculated. Chemotherapy response was assessed stratified by subtype. Radiomic feature importance was ranked.

**Results:** The mean±STDEV sensitivity, specificity and ROC-AUC were 0.90±0.07, 0.92±0.11, and 0.93±0.07, respectively. Patients with a KRT81+ subtype experienced significantly diminished median overall survival compared to KRT81-patients (7.0 vs. 22.6 months, HR 1.44, log-rank-test P=<0.001) and a significantly improved response to gemcitabine-based chemotherapy over FOLFIRINOX (10.14 vs. 3.8 months median overall survival, HR 0.85, P=0.037) compared to KRT81-patients, who responded significantly better to FOLFIRINOX over gemcitabine-based treatment (30.8 vs. 13.4 months median overall survival, HR 0.88, P=0.027).

**Conclusions:** The machine-learning based analysis of radiomic features enables the prediction of subtypes of PDAC, which are highly relevant for overall patient survival and response to chemotherapy.

## Introduction

Pancreatic ductal adenocarcinoma (PDAC) carries the worst prognosis of all tumor entities. Complete resection, often combined with an adjuvant chemotherapy regimen, remains the only curative therapy option in PDAC. In the metastatic setting, gemcitabine/nab-paclitaxel or FOLFIRINOX-based chemotherapy have been the mainstay in the treatment of PDAC (1–3). However, although both intensified treatment protocols increased response rates up to approximately 30%, a substantial number of patients does not respond or acquires resistance in a considerably short time. Pre-clinical and clinical evidence suggests differential response of specific PDAC subtypes to these treatments. Among these, a particularly aggressive subtype, termed quasi-mesenchymal, basal-like or cytokeratin 81 positive (KRT81+) (4,5) has been investigated and found to be more sensitive to gemcitabine treatment in vitro (6) and less sensitive to FOLFIRINOX in a prospective clinical trial (7). Thus, pre-therapeutic identification of specific subtypes in pancreatic cancer is urgently required to guide individual treatment decision.

So far, molecular profiling has relied on tissue biopsies, which are prone to undersampling, not least because of this entity’s morphological heterogeneity, which manifests as a heterogenic mix of tumor cell clusters, stroma and non-tumoral cell infiltrates. In addition, molecular subtyping requires high tissue quality and is both costly and time consuming, thus at current not introduced in routine patient care.

Non-invasive diffusion weighted-magnetic resonance imaging (DW-MRI, DWI), is an imaging technique which is part of the routine diagnostic work-up in many centers. It measures the random motion of water molecules and can thus quantify tissue microstructure and heterogeneity with high sensitivity (8). Radiomics, i.e. the computer-based analysis of non-perceptual image features, provides a novel tool for the evaluation of DWI beyond traditional descriptive radiology. Recent work has shown its potential in e.g. the differentiation of tumor grading or the prediction of therapy response and survival in various tumor entities including PDAC (9,10).

In the current study we developed a machine learning algorithm capable of predicting clinically relevant histopathological PDAC subtypes from pre-operative DW-MRI derived ADC maps and evaluated tumor subtype-stratified overall survival for different chemotherapy regimens.

## Materials and Methods

### Study design

The study was designed as a retrospective observational cohort study matched on histopathological tumor subtype.

Data collection, processing and analysis were approved by the institutional ethics committee (Ethics Commission of the Faculty of Medicine of the Technical University of Munich, protocol number 180/17). The requirement for consent was waived. All procedures were carried out in accordance to pertinent laws and regulations.

The STROBE checklist and inclusion flowchart can be found in the supplemental material. In brief, we considered 102 consecutive patients with final histopathological diagnosis of PDAC of the head or body for inclusion in the study. Patients without a final diagnosis of PDAC, with *unclassifiable* tumor subtype, who had undergone prior therapy (chemotherapy, resection prior to enrolment), died within the first 6 weeks of follow-up (to limit bias from postoperative complications), did not undergo the full imaging protocol or did not have technically sufficient imaging available (due to e.g. motion artifacts or stent placement), were excluded. A total of 55 patients who underwent surgical resection in curative intention were included in the study using histopathological subtype as the matching criterion. 27 patients with a KRT81+ subtype and 28 patients with a KRT81-subtype (5) were included. The follow-up interval began on the 1^st^ of January, 2010 and ended on the 31^st^ of December 2016. All patients died within the follow-up interval thus observed (uncensored) endpoint data is available for all patients. For 21 patients, follow-up data and histopathological data was sourced from the “PR2” cohort described in (5). For all other patients, clinical follow-up was handled by the departments of surgery and internal medicine, clinical data was sourced from the hospital’s clinical system and histopathological data was generated during the study. Radiomic data for all patients was generated during data analysis. All analyses were performed on pseudonymized datasets by separate individuals (G.K. and S.Z.) from January to May 2019.

### Clinical data

The following clinical data was collected: age at diagnosis, sex, pTNM, R, G, tumor volume (from the final histopathological report), ECOG-status, adjuvant chemotherapy (gemcitabine-based vs. no chemotherapy), palliative chemotherapy (gemcitabine-based vs. FOLFIRINOX) and lymph-node ratio (LNR). Overall survival was defined as the time from diagnosis to disease-related death.

### Imaging data

Patients underwent magnetic resonance imaging (MRI) at 1.5T (Siemens Magnetom Avanto, release VB17). The protocol included the following sequences: axial and coronal T2-weighted spin echo (SE) images at 5mm; axial T1w gradient echo (GE) images at 5mm before contrast media injection and during the arterial, pancreatic parenchymal, portal-venous, systemic venous and delayed phases (as determined by testing bolus injection); axial unidirectional diffusion-weighed imaging at b-values of 0, 50, 300 and 600 with echo-planar imaging (EPI) readout and ADC map calculation. ADC map reconstructions were 5.5×5.5×5 mm (xyz) to a 192×192 voxel matrix. Furthermore, single-shot T2w magnetic resonance cholangiopancreatography (MRCP) was performed and reconstructed as a radial maximum intensity projection (MIP) series. The imaging protocol, and the technical software and hardware specifications of the MRI machine remained unaltered during the data acquisition period.

### Image segmentation

The datasets were exported in pseudonymized form to a segmentation workstation running ITK-SNAP v. 3.8.0 (beta). Segmentation was performed under radiological reporting room conditions by consensus reading of two experienced observers (G.K. and S.Z.). After a period of two weeks, datasets were shuffled by a third person (F.L.) and segmented again by the same observers. Segmentations were then quality-controlled by an abdominal radiologist with >10 years of experience in pancreatic MRI (R.B) and the best segmentations retained. Segmentation was performed manually in the b=600 images and transferred to the ADC maps. All other sequences were available to observers for anatomical correlation.

### Biostatistical and machine learning modeling

For assessing bias due to clinical confounders, overall survival time was evaluated by a multivariate *Cox proportional hazards* model. The distributions of covariates were compared between groups with different histopathological subtype using *Fisher’s exact test*.

Biostatistical modeling was performed using the Python (v.3.7.3) package *Lifelines.* Kaplan-Meier-Plots were drawn in GraphPad Prism (v.8). For all inferential statistical procedures, a P-value of <0.05 was considered statistically significant.

Image postprocessing, feature extraction, feature preprocessing, feature engineering and machine learning modeling are described in the supplemental material. In brief, radiomic features were derived using *PyRadiomics* (v. 2.1) (11) yielding a total of 1606 features, of which 40 were retained after exclusion of features with low-variance or repeated segmentation instability. A supervised *Gradient Boosted Decision Tree* model (*XGBoost* (12), instantiated as a binary classifier within the Python library *scikit-learn*) was fit with histopathological subtype as a binary label to the radiomic features and tested for predictive sensitivity, specificity and ROC-AUC. Training was performed by randomized 10-fold shuffle-splitting cross-validation on 70% (N=38) of the cohort and the model was tested on the remaining-unseen-30% (N=17) of the cohort. Significance testing for model evaluation metrics was carried out using permutation testing (13). The threshold probability for classification the default value of. 50. Feature importance was assessed by the inbuilt feature importance classifier (using the “gain” parameter).

### Histopathological workup

Histopathological staining and immunohistochemical workup were performed as described in (5) and tumors were categorized into either one of two classes: KRT81+ or KRT81-(Fig 1).

**Fig 1.**
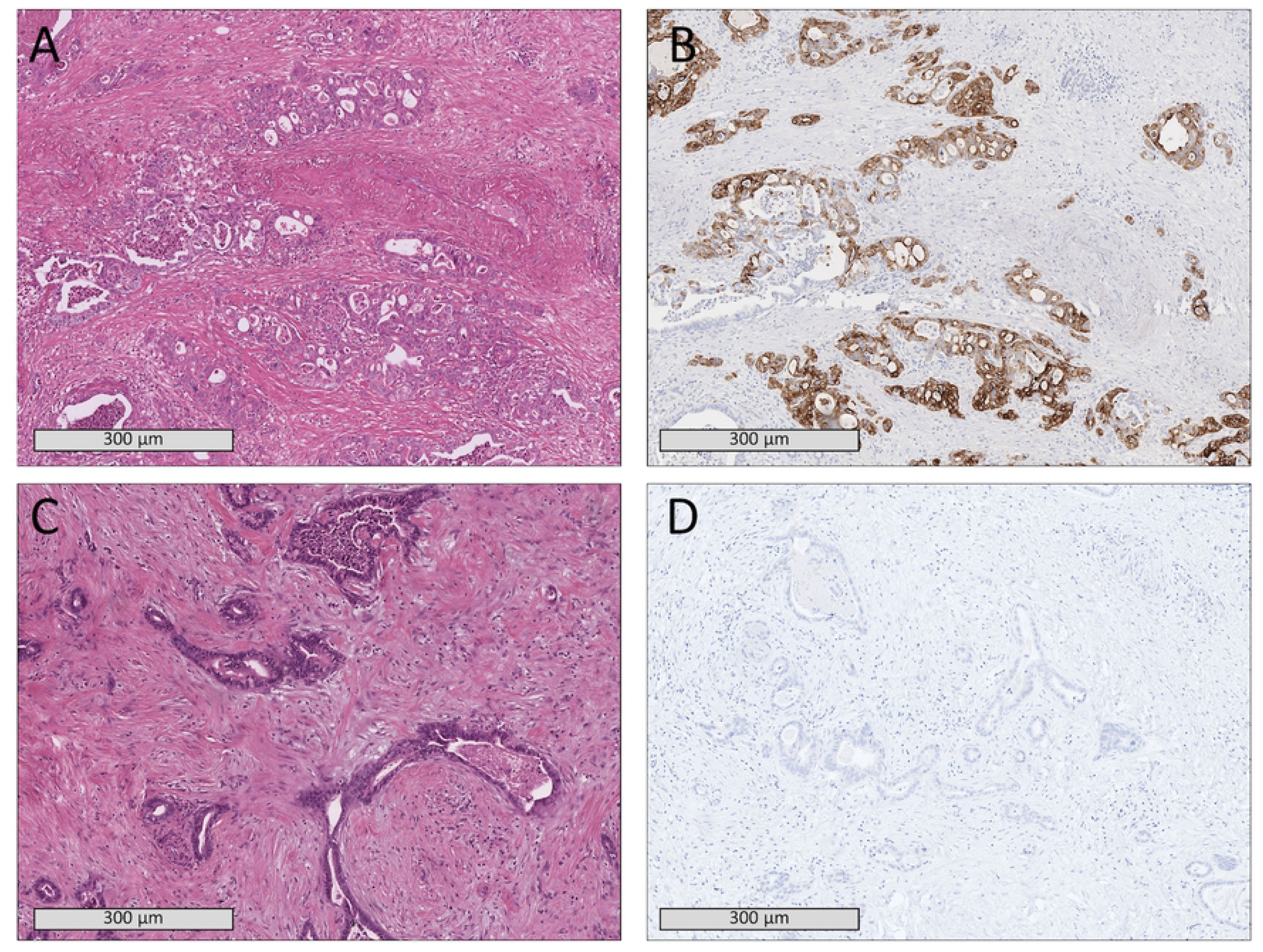
Histopathological samples of two patients showing comparable tissue morphology in H&E staining (A,C) but a KRT81+ subtype (B) in one patient and KRT81-subtype (D) in the other patient.

## Results

The molecular subtype of PDAC was significantly associated with overall survival. Patients with a KRT81+ subtype experienced significantly diminished overall survival (7.0 [1.93 to 29.0] vs. 22.6 [2.63 to 96.97] months median survival, HR 1.44 [0.76-2.12], log-rank-test P=<0.001, Fig 2, Table 1). No other covariate was significantly associated with overall survival in this cohort and the baseline distribution of clinical covariates did not differ significantly between the two patient subcohorts (Table 2).

**Table 1.**
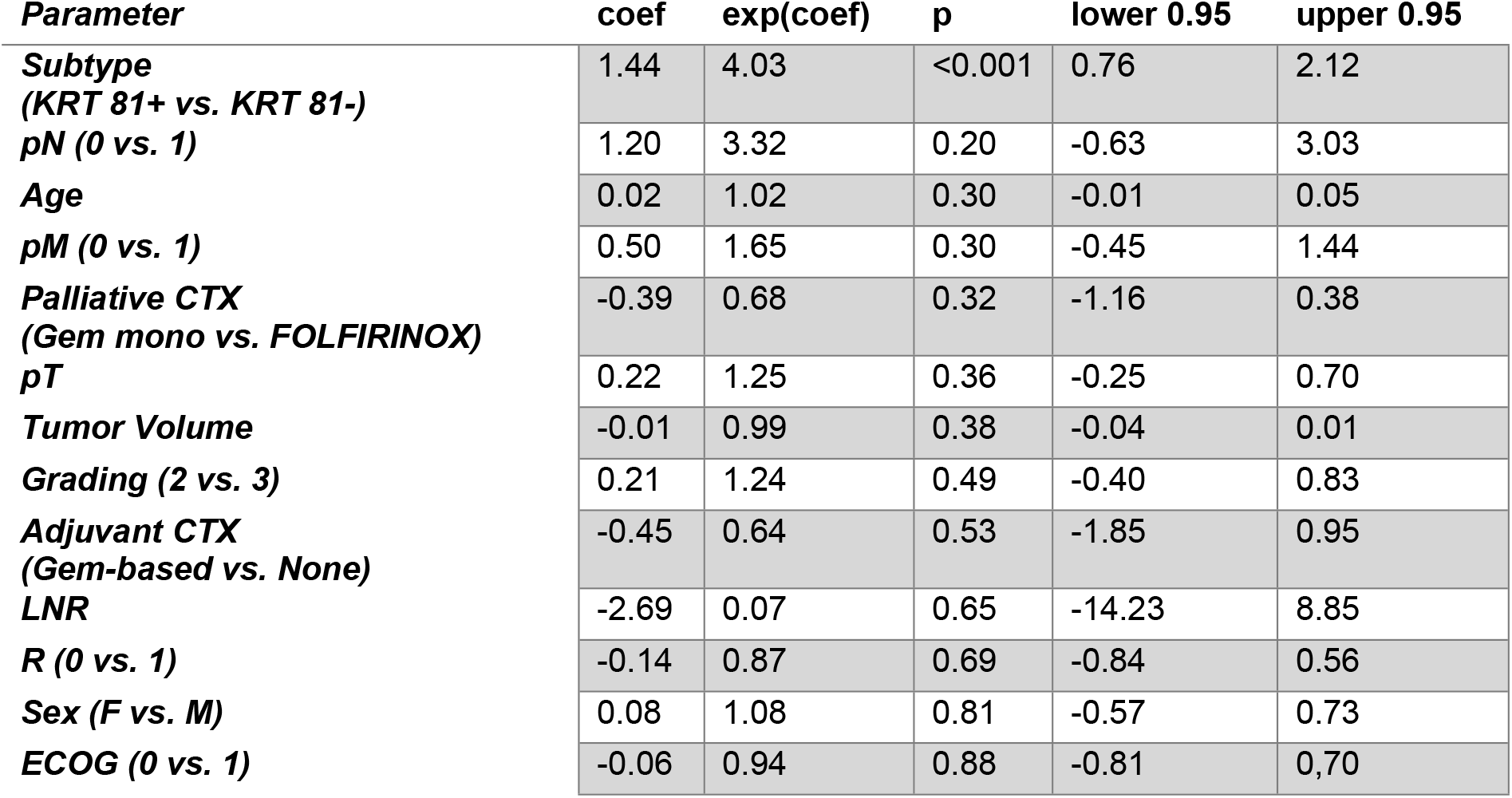
Cox proportional hazards analysis results of clinical parameters.

| *Parameter* | **coef** | **exp(coef)** | **p** | **lower 0.95** | **upper 0.95** |
| --- | --- | --- | --- | --- | --- |
| **Subtype**<br>*(KRT 81+ vs. KRT 81-)* | 1.44 | 4.03 | <0.001 | 0.76 | 2.12 |
| **pN** (0 vs. 1) | 1.20 | 3.32 | 0.20 | -0.63 | 3.03 |
| **Age** | 0.02 | 1.02 | 0.30 | -0.01 | 0.05 |
| **pM** (0 vs. 1) | 0.50 | 1.65 | 0.30 | -0.45 | 1.44 |
| **Palliative CTX**<br>*(Gem mono vs. FOLFIRINOX)* | -0.39 | 0.68 | 0.32 | -1.16 | 0.38 |
| **pT** | 0.22 | 1.25 | 0.36 | -0.25 | 0.70 |
| **Tumor Volume** | -0.01 | 0.99 | 0.38 | -0.04 | 0.01 |
| **Grading** (2 vs. 3) | 0.21 | 1.24 | 0.49 | -0.40 | 0.83 |
| **Adjuvant CTX**<br>*(Gem-based vs. None)* | -0.45 | 0.64 | 0.53 | -1.85 | 0.95 |
| **LNR** | -2.69 | 0.07 | 0.65 | -14.23 | 8.85 |
| **R** (0 vs. 1) | -0.14 | 0.87 | 0.69 | -0.84 | 0.56 |
| **Sex** (F vs. M) | 0.08 | 1.08 | 0.81 | -0.57 | 0.73 |
| **ECOG** (0 vs. 1) | -0.06 | 0.94 | 0.88 | -0.81 | 0.70 |

**Table 2.**
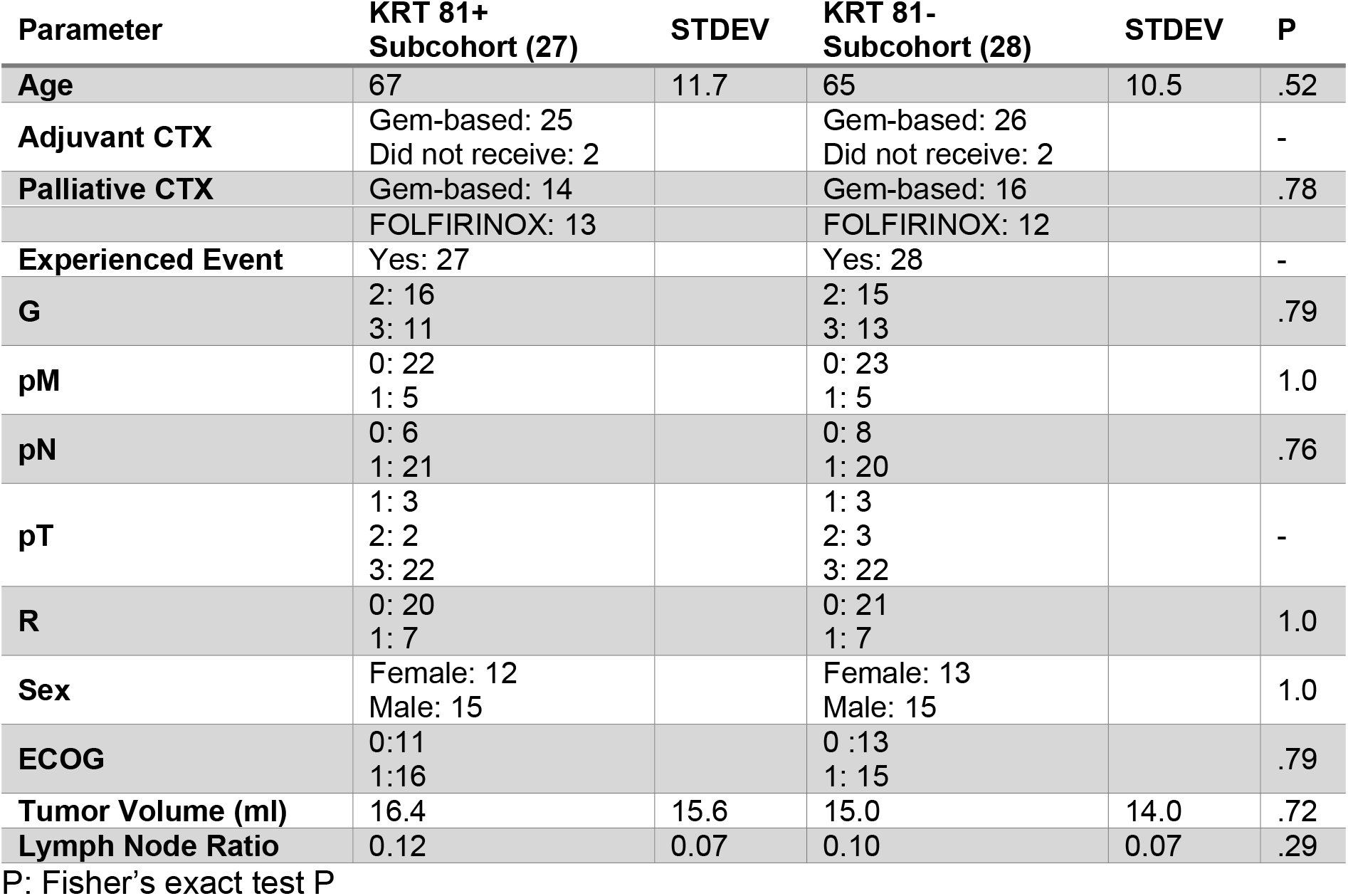
Distribution of clinical parameters between the cohorts with KRT81+ and KRT81-tumor subtypes alongside crosstabulation results

| **Parameter** | **KRT 81+ Subcohort (27)** | **STDEV** | **KRT 81- Subcohort (28)** | **STDEV** | **P** |
| --- | --- | --- | --- | --- | --- |
| **Age** | 67 | 11.7 | 65 | 10.5 | .52 |
| **Adjuvant CTX** | Gem-based: 25<br>Did not receive: 2 |  | Gem-based: 26<br>Did not receive: 2 |  | - |
| **Palliative CTX** | Gem-based: 14 |  | Gem-based: 16 |  | .78 |

|  | FOLFIRINOX: 13 |  | FOLFIRINOX: 12 |  |  |
| --- | --- | --- | --- | --- | --- |
| **Experienced Event** | Yes: 27 |  | Yes: 28 |  | - |
| **G** | 2: 16<br>3: 11 |  | 2: 15<br>3: 13 |  | .79 |
| **pM** | 0: 22<br>1: 5 |  | 0: 23<br>1: 5 |  | 1.0 |
| **pN** | 0: 6<br>1: 21 |  | 0: 8<br>1: 20 |  | .76 |
| **pT** | 1: 3<br>2: 2<br>3: 22 |  | 1: 3<br>2: 3<br>3: 22 |  | - |
| **R** | 0: 20<br>1: 7 |  | 0: 21<br>1: 7 |  | 1.0 |
| **Sex** | Female: 12<br>Male: 15 |  | Female: 13<br>Male: 15 |  | 1.0 |
| **ECOG** | 0:11<br>1:16 |  | 0 :13<br>1: 15 |  | .79 |
| **Tumor Volume (ml)** | 16.4 | 15.6 | 15.0 | 14.0 | .72 |
| **Lymph Node Ratio** | 0.12 | 0.07 | 0.10 | 0.07 | .29 |

P: Fisher's exact test P

**Fig 2.**
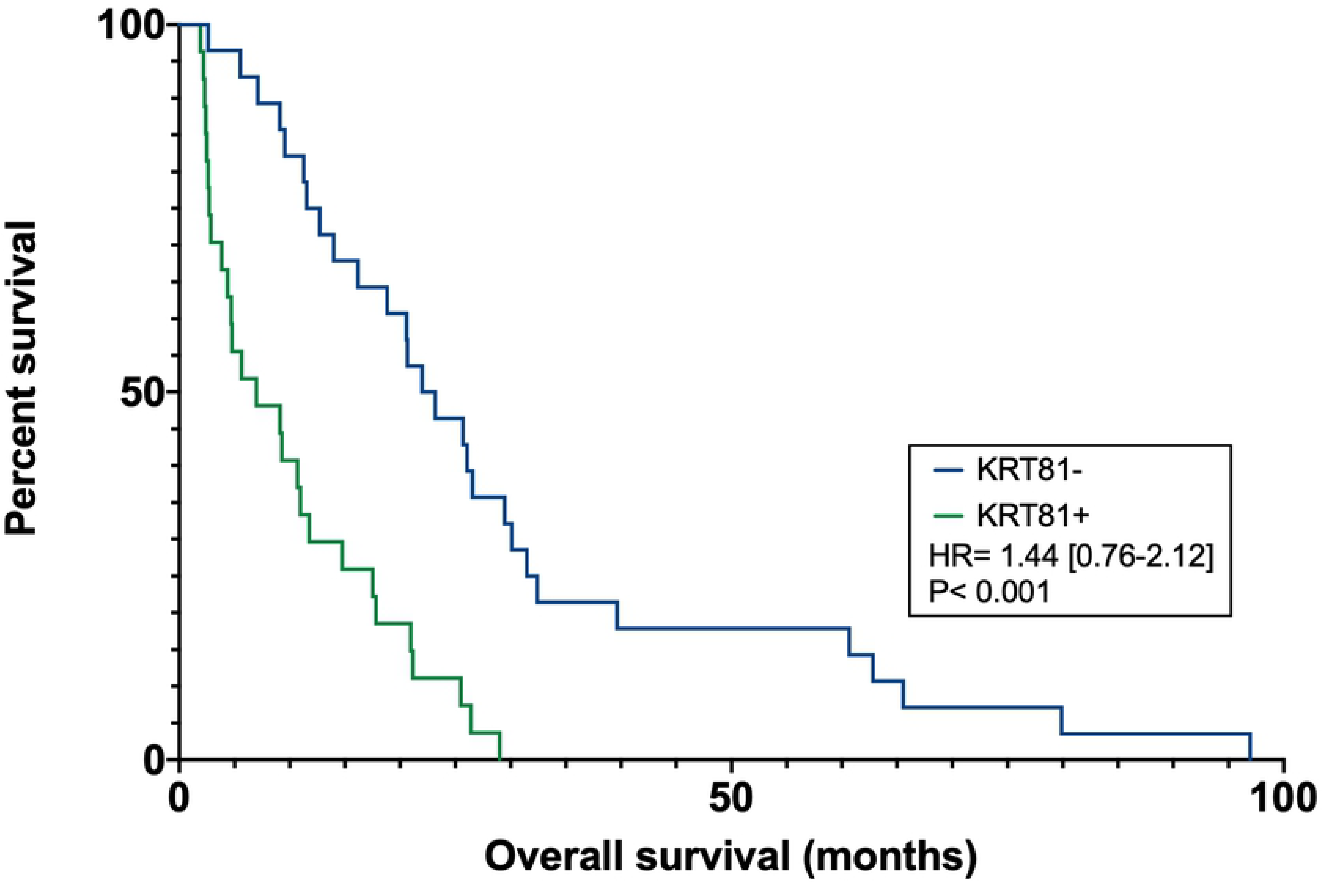
Patients with a KRT81+ subtype experienced significantly diminished overall survival.

The machine learning algorithm achieved a mean±STDEV sensitivity, specificity and ROC-AUC of 0.90±0.07, 0.92±0.11, and 0.93±0.07, respectively; all P=0.01 (Fig 3).

**Fig 3:**
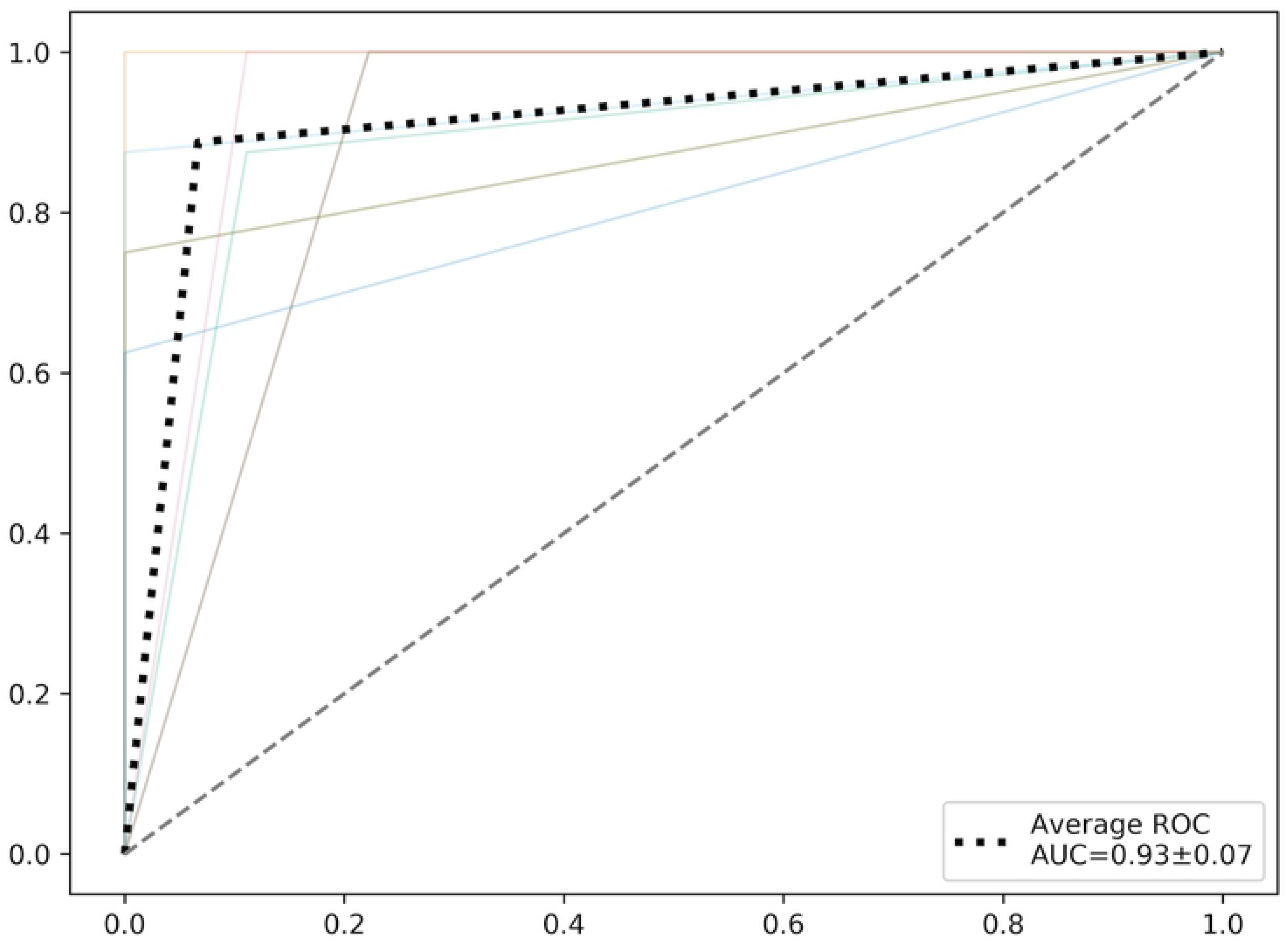
ROC curves (colored) and average ROC-curve (black dotted) over 10 random stratified shuffle-splits of the dataset.

The feature importance evaluation of the algorithm yielded 13 radiomic parameters with an importance greater than zero for the classification process. Among these, entropy derived from the original image was classified as the most important feature by a large margin. All features alongside their importance metrics can be found in Table 3.

**Table 3:**
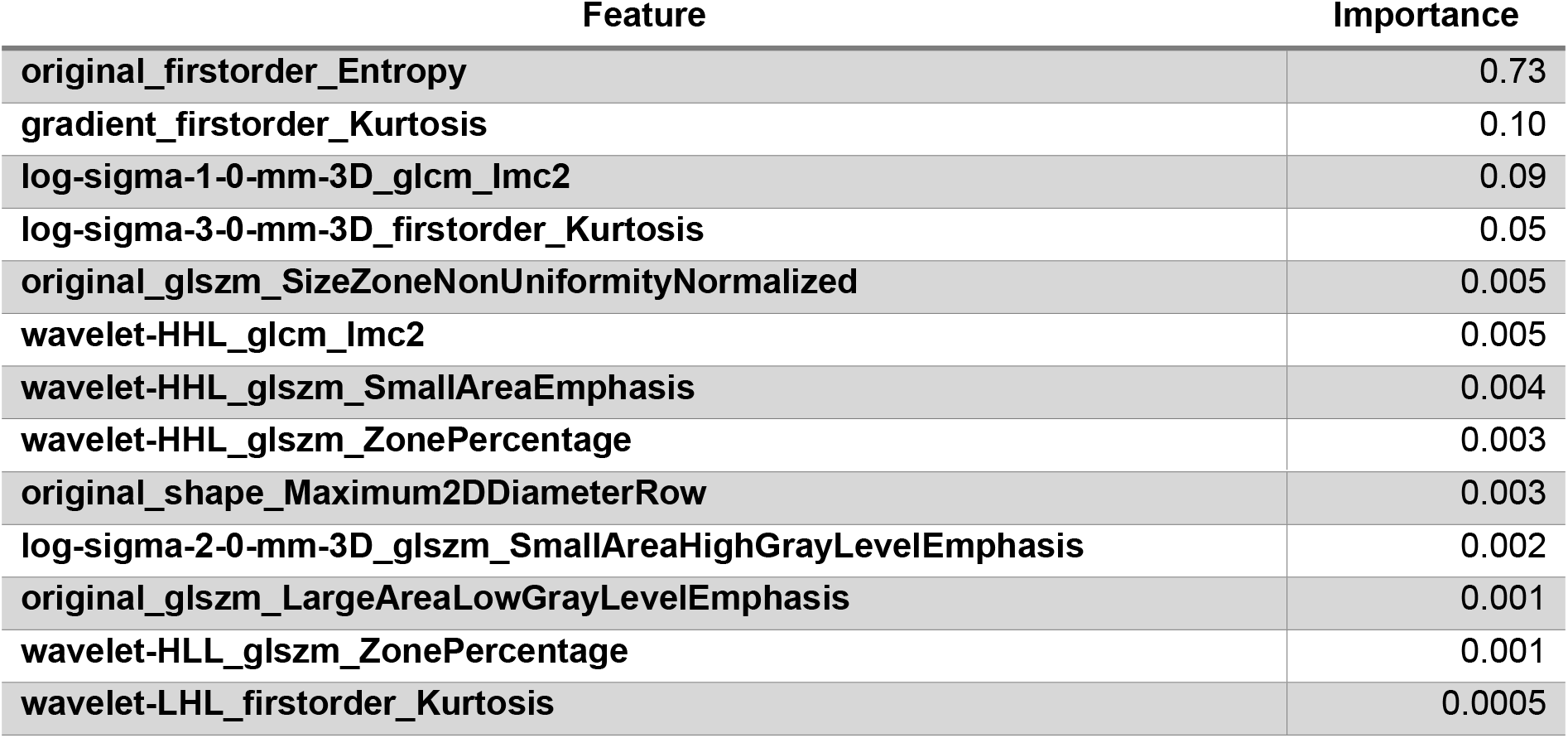
Radiomic features alongside their importance as ranked by the algorithm

Overall survival was evaluated separately for histopathological subtypes stratified by chemotherapy regimen. Patients with a KRT81+ histopathological subtype who received gemcitabine-based palliative chemotherapy experienced significantly improved survival compared to patients with KRT81+ tumors who received FOLFIRINOX (10.14 vs. 3.8 months median survival, HR 0.85 [0.02-1.67], P=0.037, Fig 4). Conversely, KRT81-subtype patients experienced significantly improved survival under FOLFIRINOX chemotherapy compared to gemcitabine-based regimens (30.8 vs. 13.4 months median survival, HR 0.88 0.08-1.67], P=0.027, Fig 5).

**Fig 4.**
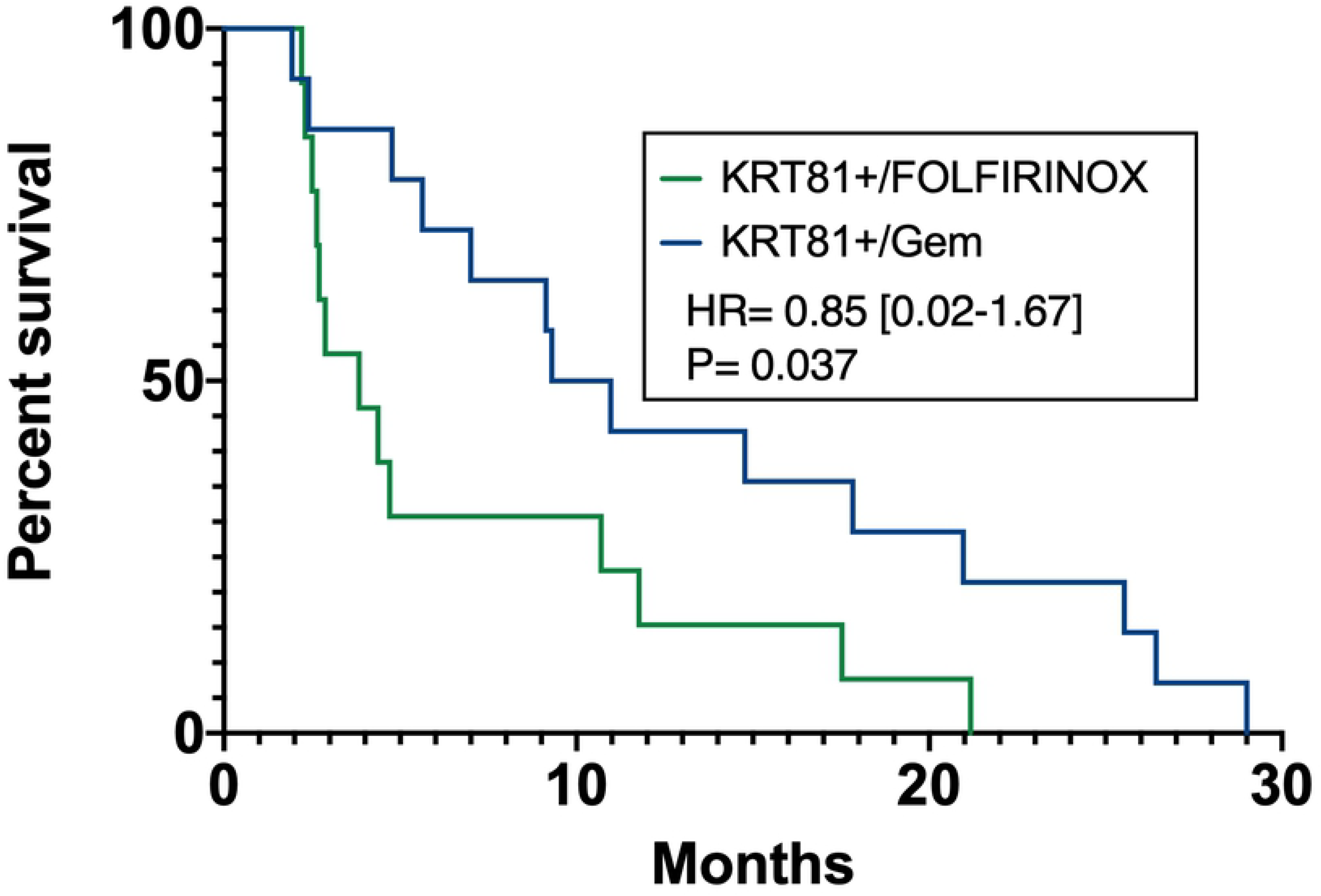
Patients with KRT81+ subtype experience longer overall survival under palliative gemcitabine chemotherapy

**Fig 5.**
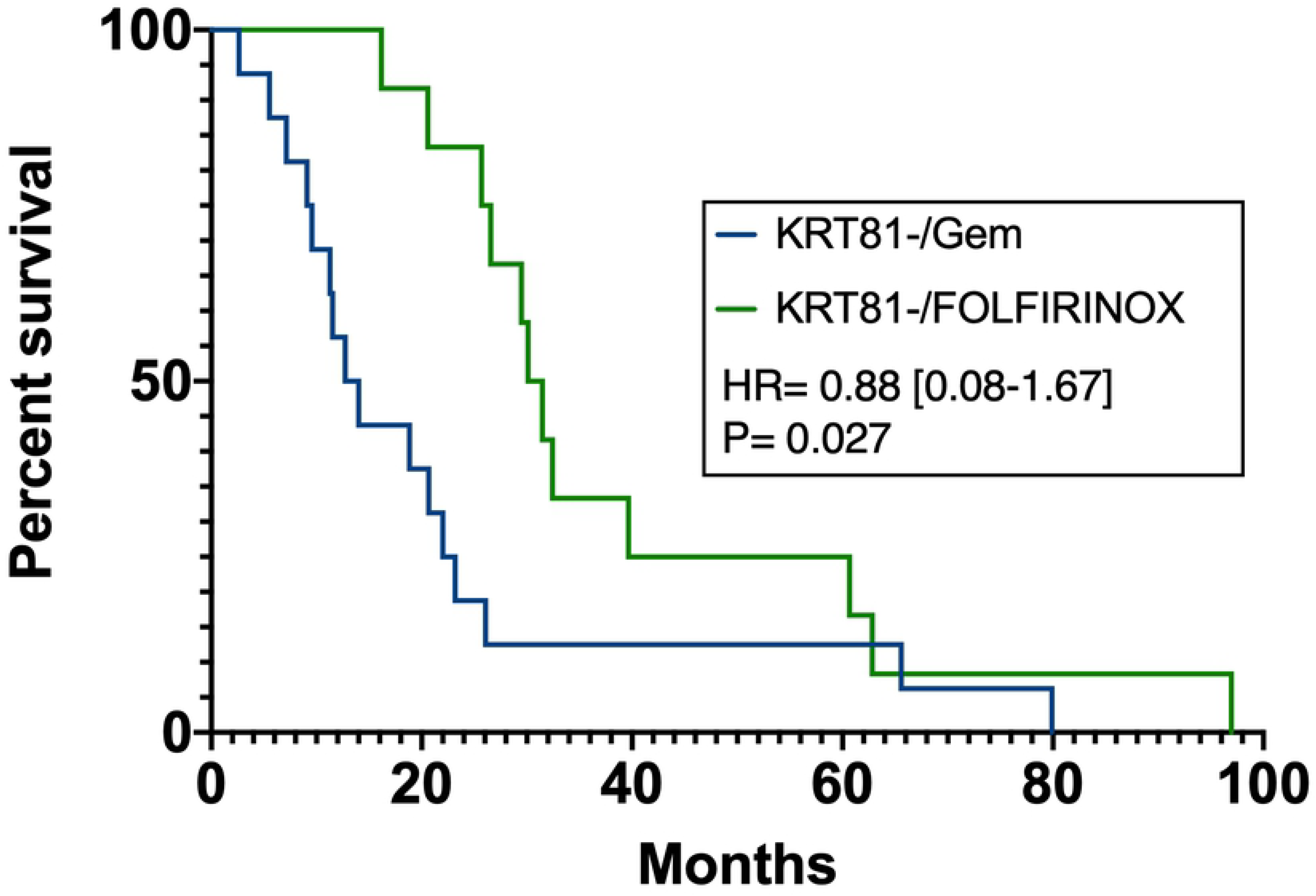
Patients with KRT81-subtype experience improved survival under palliative FOLFIRINOX chemotherapy

## Discussion

In this exploratory study, we demonstrate that radiomic analysis of ADC maps paired with machine-learning modeling can discriminate with high sensitivity and specificity between two groups of histomorphologically defined molecular subtypes of pancreatic ductal adenocarcinoma (PDAC), associated with significantly different responses to commonly employed chemotherapeutic regimens in the palliative setting. We thus provide evidence for the utility of radiomics and machine learning for the non-invasive stratification of pancreatic cancer patients.

The potential of non-invasive imaging-derived biomarkers (from non-perceptual image features or source data) has been demonstrated in several studies with the prediction of tumor genetics and patient outcome (14–16). However, their widespread application beyond proof-of-principle studies requires the identification of stable and reproducible parameters, embedded within a standardized and quality-controlled workflow (17–20).

Among the parameters tested for classification in our study, entropy was ranked the most important by the algorithm. This finding is encouraging, since entropy and entropy-related features, which express disorder and heterogeneity of the image, have been demonstrated in (meta-)analyses of different tumor entities, across modalities as promising candidate parameters (21–23). We found whole-tumor entropy values highly predictive of the basal-like PDAC subtype. However, considering immanent sampling errors in this histopathologically heterogeneous tumor entity, the complexity of mutational events (e.g. variable amounts of mutational *Kras* (24) and the likelihood of ongoing transitional processes, entropy as a continuous variable can be imagined as a non-invasive “sensor” of such events that correlates with the extent of basal-like regions within a particular tumor. The testing of such hypotheses is challenging, requiring an integrated whole-tumor analysis, including high data-rich imaging, histopathology and molecular profiling (25).

The rapid evolution of new therapeutic options in the treatment of PDAC requires the development of markers for a reliable pre-therapeutic patient stratification and -in light of the above-mentioned plasticity, therapy monitoring. Conroy et al. demonstrated significantly improved survival rates of FOLFIRINOX over Gemcitabine monotherapy in the palliative setting (3). However, the COMPASS trial (7) demonstrated differential response of the basal-like versus non-basal-like PDAC subtypes to FOLFIRINOX treatment, which is well in accordance with our study results. If further validated in prospective trials, these findings could have tremendous implications in patient stratification and subtype-guided therapy selection. In addition, targeted therapies such as Olaparib, are highly effective yet even more specific for a certain molecular profile (26) and many new targeted, stroma- and immune-based treatment strategies are being explored. This increasing complexity requires robust and cost-efficient tools for clinically relevant patient stratification to best leverage current knowledge and advance the field. Informed decision based on molecular profiling (microdissection and genome sequencing) as applied in the COMPASS trial faces serious limitations (i.e. sampling error, high cost) and is therefore currently not feasible in routine patient care. Quantitative noninvasive imaging, and especially radiation and contrast-free quantitative modalities such as DWI may serve this purpose and are thus excellent candidates for exploration in a prospective trial design.

Limitations of this study are the small cohort size and lack of an external testing cohort as well as the retrospective, single-center nature of the investigation. Such issues are still common in the imaging field and compounded by the lack of standardization in sequence acquisition between institutions and of overarching registers or study centers permitting patient pooling. Recently, initiatives have arisen to combat some of these issues by harmonization of MRI protocols (27) and the standardization of imaging markers (28).

In conclusion, our study is an exploratory venture into the field of quantitative imaging analysis and radiology/pathology-correlation in PDAC. We encourage the validation of our findings in a larger cohort and in a prospective trial design.

## Acknowledgments

Authors wish to thank Irina Heid for the ongoing support.

